# Potential *L-asparaginase* producing *E.coli* sources among River water and cafeteria sewage: The case of Wabe River and Wolkite University students’ cafeteria in Wolkite, Ethiopia

**DOI:** 10.1101/2024.05.23.595502

**Authors:** Kajelcha Fikadu Tufa, Senbeto Beri Mosisa, Aklilu FikaduTufa, Admas Berhanu Beriso

## Abstract

*L-asparaginase* is a promising enzyme for cancer treatment and is found in plants, animals and microbes. This enzyme is of great medical and industrial importance. It is used with inside the remedy of acute lymphoblastic leukemia and helps in reducing the acryl amide substances found in fried and baked foods. Its source varies from bacteria to yeast and fungi. This study aimed to screen the potential *L-asparaginase* producing E. coli isolates among river water and cafeteria sewage samples near the Gubryie area, SNNPR-Ethiopia. In this study, *E. coli* isolates were isolated from sewage from the Wabe River and Wolkite University student cafeteria. During the study, 14 isolates, 11 from cafeteria sewage and 3 from Wabe River, were confirmed to be *E. coli* using IMViC, TSI, SCA, and Gram tests. For the *E. coli*-positive samples, screening of *L-asparaginase* was performed using the phenol red indicator. The change in color from yellow to pink in M9 media due to the acidic environment created when *L-asparagine* was degraded to urea indicates the presence of *L-asparginase* in the potent *E. coli* cells. The production of *L-asparaginase* was carried out using submerged fermentation method. Mechanical cell disruption method, high speed centrifugation, was used to separate the secreted enzyme from cells. The potential of the *E. coli* cells to produce *L-asparginase* was also checked using a rapid plate assay method with the indicator dye phenol red. The zone of inhibition for the intracellular enzyme activity ranges from 16.5 mm up to 22.25 mm while that of extracellular enzyme ranged from 7.5 mm up to 9 mm. The commonly used software system is SPSS version 23. The PCR result depicted that the *ansA* gene presence was confirmed in 50% of the isolates. The result confirmed that the *E*.*coli* isolates from sewage showed better *L-asparaginase* production potency than the Wabe River isolates. This study indicated that *E. coli* strains are promising sources of *L-Asparginase* for food and pharmacological companies if the scale-up of this work has been completed in the future.

## 1. Introduction

*L-asparaginase* is an enzyme that acts on the amino acid *L-asparagine* and is mostly used as an antitumor agent. It contributes one-third of the global demand as an antileukemic and antilymphoma agent, and it is mainly used to treat ALL in combination with vincristine and a glucocorticoid (Abdelrazek *et al*., 2020). There are two types of L-ANases namely extracellular and intracellular L-ANases. They are naturally available enzymes expressed and produced by animal tissues, bacteria, plants, and the serum of certain rodents but not by humans (Castro *et al*., 2021).

L-Asparagine is a very important amino acid that is used as a nutritional source for both healthy cells and tumor cells. Cancer cells are dependent on an exogenous supply of L-asparagine from body fluids since they cannot synthesize amino acids. As several types of tumor cells require L-asparagine, which is an essential amino acid for protein synthesis, they are deprived of an essential growth factor in the presence of L-ANase. Effective depletion of L-asparagine results in cytotoxicity toward leukemic cells. The administration of L-ANase within the body does not affect the function of normal cells because L-ANPases inherently synthesize L-asparagine according to their own requirements but reduce its concentration in the plasma pool (Castro *et al*., 2021).

The purpose of this motive is that it is miles biodegradable and nonpoisonous and may be easily administered on neighborhood webpages. Among the different sources of L-ANase producers, the principal sources of these enzymes are from the microbes *Escherichia coli* and Erwiniaspp, which are currently in clinical use as drugs (business brands: erwinase and colpase) within the control of lymphoblastic leukemia due to their excessive substrate affinity (Vaishnavi and Palaniswamy, 2023).

*Escherichia coli* are a gram-negative, rod-fashioned bacterium that is generally found within the decreased gut of warm-blooded organisms (endotherms). Most *E. coli* traces are harmless; however, a few serotypes can cause critical meal poisoning in humans. The innocent traces are a part of the ordinary flowers of the gut and might gain their hosts through generating diet K2 and through stopping the established order of pathogenic microorganisms in the intestine. EMB agar media is the most common medium for isolating *E. coli* because it is selective for gram-negative bacteria, especially *E. coli* (‘Key facts’, 2018; Acharya Tankeshwar, 2023).

## 2. Materials and Methods

### 2.1 Sample collection and transportation

Samples were collected from and around the Gubriye area, i.e., from the Wabe River and sewage from the cafeteria at Wolkite University. The samples were transported to Molecular Biotechnology laboratory at Wolkite University and stored at 4°C until the work start.

## 2.2 Isolation and identification of *E. coli*

Here, the colony number, color differentiation, and diameter of the zone of colony were estimated. Under aseptic conditions, 7 sterile test tubes were labeled 10-1, 10-2, 10-3, 10-4, 10-5, 10-6 and 10-7. Nine millilitres of distilled water was added to each test tube. One milliliter of sample was added to the first tube, after which serial dilution was performed. From each tube, 1 ml was inoculated on Eosin Methylene Blue agar. The plates were then incubated for 24 h at 37°C. The colonies were then isolated using the spread plate method on EMB (Now, 2024). The suspected colonies were selected and streaked on EMB agar plates once again using four-way streak methods for further purification, after which the colonies were inoculated on a TSI slant and citrate slant after purification. The next day, the isolates were further streaked onto EMB agar plates to maintain a pure culture of bacteria that was used to screen for L-ANase production.

### 2.3 Biochemical characterization

A series of biochemical tests, including the IMViC, TSI, and catalase tests, were performed on the presumptive colonies according to the protocol reported by (Gopireddy, 2011). All the tests were positive except for the citrate and Vonoges–Proskauer tests. The biochemically confirmed *E. coli* isolates were then subjected to a screening step. The isolated strains were screened for potential L-ANase production. To do this, two modified nutrient agar plates were taken and divided into sectors. One plate was left undisturbed and served as a quality control. Then, all the plates were kept in an incubator.

Pure colonies of the isolates were tested for Gram-stained bacteria, and the results are shown in pink under a microscope after treatment with the sample Gram-conjugated chemicals. Initially, both garlic-positive and gram-negative bacteria develop violet colors. However, after treatment of the colonies with iodine and safranin, the gram-negative bacteria developed a pinkish color. After Gram staiing characterization, a series of biochemical tests were used for characterization of the isolates. Then, a Simmon’s citrate agar test was performed to determine whether the microorganisms were able to utilize citrate as a carbon source. All the isolates were negative according to the Simon citrate test. This means that E. coli was unable to use citrate as a carbon and nitrogen source. All isolates were subjected to IMViC tests which include Indole test, Methyl Red test, Voges Proskauer test, and Catalase test. Among these tests, only Voges Proskauer test resulted negative for the isolates which depicted the presence of *E*.*coli* isolates.

### 2.4 Production of Extracellular L-ANase

Isolates that showed good potential during screening were streaked on Eosin methylene blue agar, and the same isolate(s) were then streaked on M9 media and non-L-asparagine-containing media (containing only phenol red) to serve as a negative control. Plates were kept in an incubator. M9 medium was used for differentiating L-ANase-producing bacterial species from nonproducer strains. On the next day, pure cultures of the isolates were inoculated into 5 ml of nutrient broth using a sterile wire loop and incubated at 37°C for 24 hrs in an incubator. For the production of L-ANase, production media were prepared in a 250 ml Erlenmeyer flask containing 0.5% NaCl (5 g/L), maltose sugar (10 g/L), KH2PO4 (0.75 g g/L), L-aspartic acid (100 g/L), and 0.009% (v/v) phenol red as an indicator. The pH was maintained at 7.4, and the solution was autoclaved. Using a micropipette, medium was added to the cuvette, and the optical density was determined. After that, 1 ml of the overnight bacterial broth (O.D>0.5) was inoculated in 100 ml of production media and incubated in a horizontal shaker at 120 rpm for 3 days.

The optical density (OD) was then measured using a UV spectrophotometer at 460 nm. That is, the OD of the growth in the medium. Note the pH of the production medium. A low pH confirmed the enzyme production. In an aseptic environment, take appropriate quantity from production media using sterile tip of micropipette and add it into Eppendorf’s tube after measuring the OD and PH of each sample and centrifuge it using a micro centrifuge machine (Bio maker). The pellet and the supernatant were separated. The supernatant was obtained from a crude enzyme extract (extracellular enzyme), while the pellet contained cell debris. The supernatant was transferred to a sterile cuvette, and the O.D. was measured at 460 nm. All the supernatants were carefully collected in sterile test tubes using sterile micropettings, after which the resulting mixture was marked as the extracellular enzyme extract. Put a cotton plug on it, and preserve this in a freezer until use (in a sterile condition) (Sharma, Verma and Pandey, 2019).

### 2.5 Enzyme Assay

A plate assay was used to detect L-ANase activity on modified nutrient agar supplemented with beef extract, peptone extract, L-asparagine, sodium chloride, and phenol red (the pH of this medium was >6.8, at which point the phenol red turned orange). The reason for keeping the medium orange was that it was helpful for reading and measuring the red zone in contrast to the orange color of the agar. A control agar sample was also used in this assay to serve as a negative control. Using the sterile tip of the micropipette, a drop of an extracellular enzyme was added to the wells of both plates, and both plates were incubated. The pink zone around the holes indicates enzyme activity because the enzyme causes the acryl amide and/or L-aspartic acid to break down, and ammonia is released, which increases the pH (alkaline); additionally, the phenol red present in the medium turns pinkish red. Zones were measured in mm (Mahajan *et al*., 2013).

### 2.6. PCR amplification of potential *L-Asparaginase* producing *E*.*coli* Isolates

Genomic DNA was extracted from the bacterial isolates using boiling standard methods. To do this, 2 mL of the overnight E. coli culture was added in to Eppendorf tube and centrifuged to pellet the cells. The supernatant was discarded and the cell pellet was resuspended in 100 µL of nuclease free water. The cell suspension was then boiled at 100°C for 10 minutes to lyse the cells followed by cooling the boiled suspension to room temperature for more than 30 minutes. The suspension was thawed and centrifuged at 1400rpm for 5minutes to pellet the cell debris. Transfer the supernatant containing genomic DNA to a new sterile Eppendorf tube. The DNA extract was stored at -20°C until used for PCR.

Polymerase chain reaction machine is a very important tool of molecular biology for molecular characterization of an organism under study. In this study, specific primers ((*ansA*-F): 5’-ATGCGTGCTGATTGAAAG-3’ (ansA-R): 5’-TTAACCCGCTTTTTCAC-3) for the target *ansA* gene, which encode L-asparaginase in *E. coli*. This gene was targeted because it is responsible for the production of L-asparaginase. The PCR reaction mixture includes the extracted DNA, specific primers for the target gene, dNTPs, Taq polymerase, buffer, and MgCl_2_ was prepared based on the manufacturer’s guideline. The PCR conditions were set, denaturation: 94°C for 30 seconds, annealing: 54°C for 30 seconds, extension: 72°C for 1 minute per kb of the expected product length. The PCR products were separated on an agarose gel-electrophoresis and visualized under UV light to confirm the presence of the expected size bands, indicating successful amplification of the target gene. This approach ensured the accurate identification of *L-Asparaginase* producing *E. coli* isolates, which was crucial for applications in biotechnology and medicine, particularly in the development of cancer therapeutics (Sudhir *et al*., 2012).

## 3. Results

### 3.1 Isolation of *Escherichia coli*

A total of 14 isolates were isolated and identified as a presumptive *E*.*coli* isolates from the sample sites, cafeteria sewage and river water. Among these, *most of the isolates*, 11(78.6*%)*, were from Wolkite University students’ cafeteria sewage sample while the rest 3(21.4%) were from Wabe river sample (**Figure 1**). On the next day, all suspected colonies that developed with a green metallic sheen during isolation were properly streaked on EMB media by dividing them into 4 sectors, after which a clear green metallic sheen developed (**Figure 1**).

### 3.2 Screening for *L-asparaginase* Production

In this study, biochemically confirmed *E. coli* isolates were screened on M9 media for the detection of *L-ANase* production by which L-aspartic acid was produced. Asparagine was used as a substrate, and the isolates that exhibited excellent L-ANase activity, such as those that formed a pinkish zone indicating hydrolysis of L-aspartic acid after 48 hours while the negative control was remain colorless (**Figure 2**).

**Figure 2:**
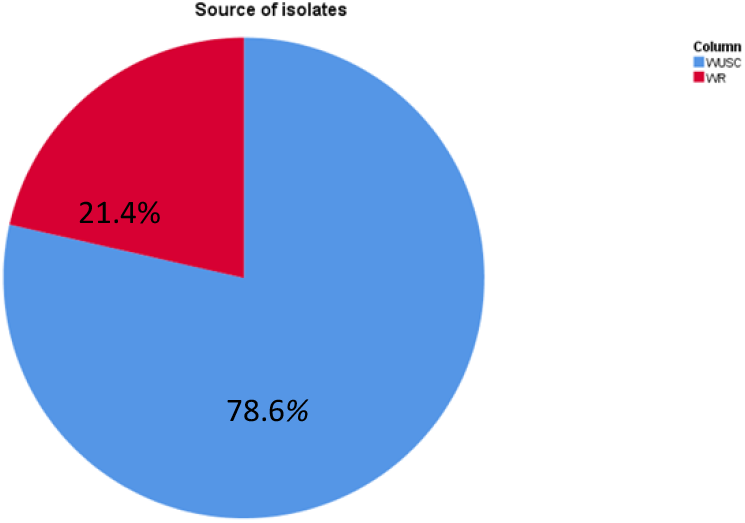
The comparision pichart: This pi-chart summarized that the number of potential *L-asparaginase* producing *E*.*coli* isolates from Wolkite University Students’ cafeteria sewage was greater than that of Wabe river water sample.

**Figure 3:**
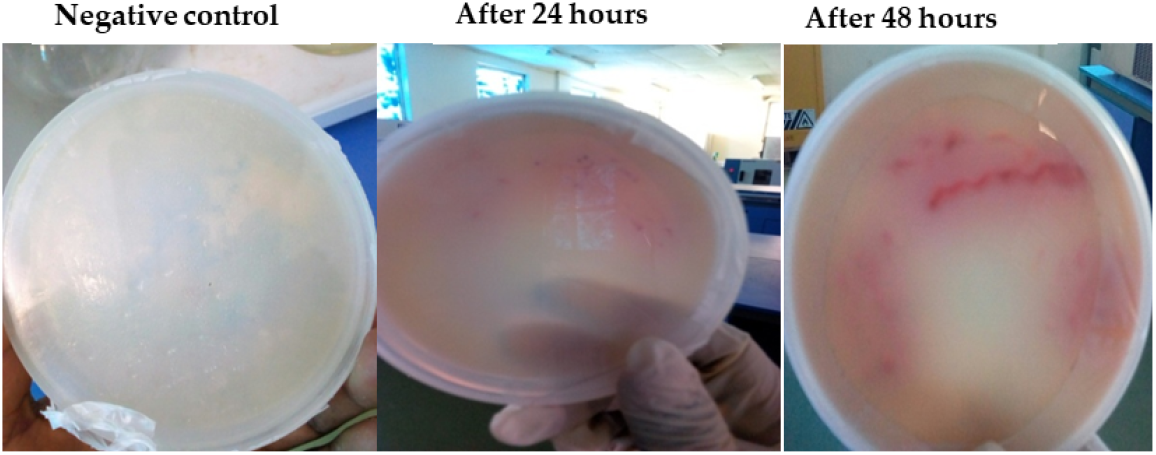
Screening the isolates for *L-asparaginase* production: Results of screening. The one-day incubation period was less effective than the two-day incubation period. Thus, after 48 hours, the enzyme was effectively incubated with the cells.

**Figure 4:**
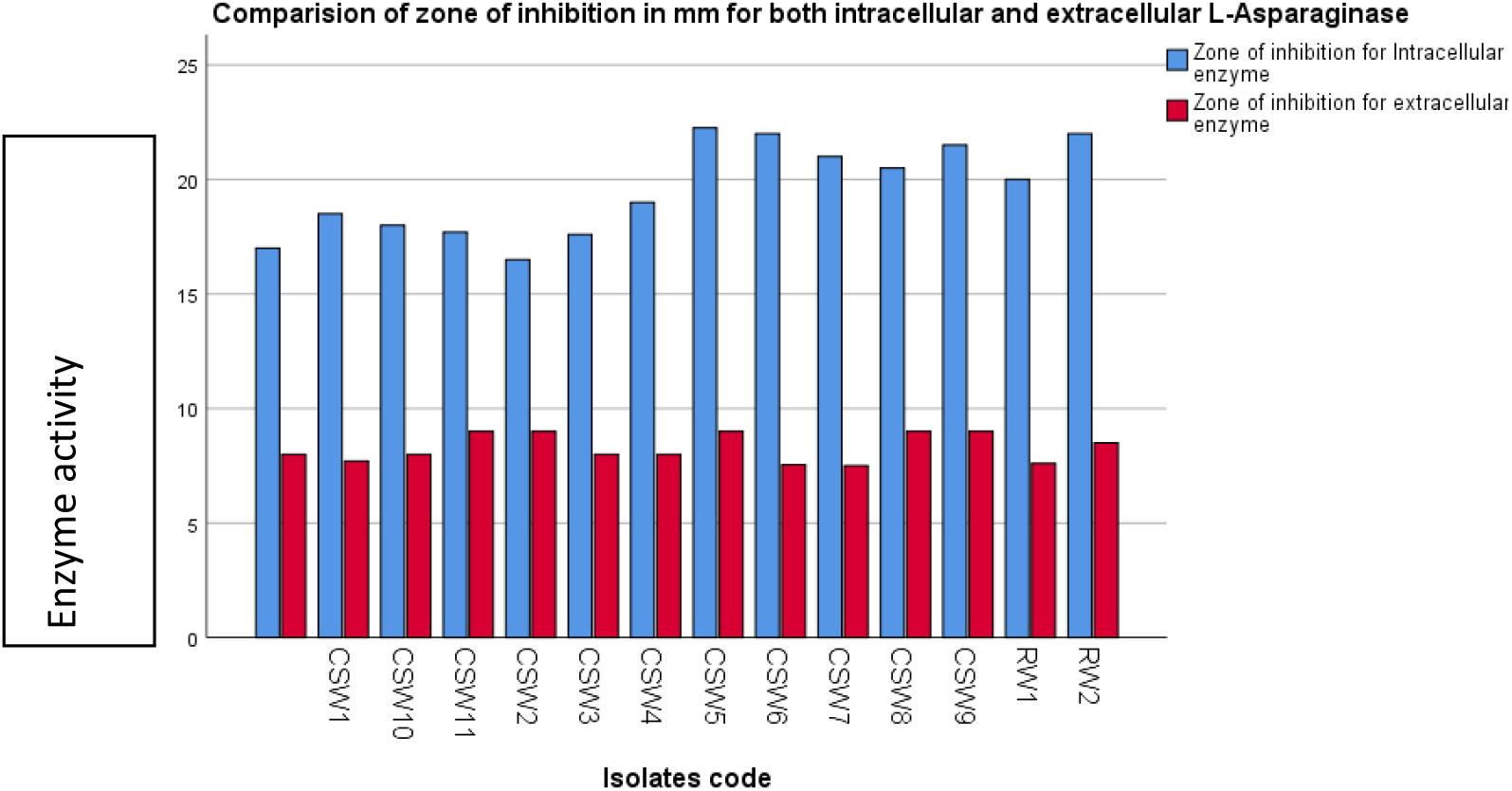
Zone of inhibition for both intracellular and extracellular *L-asparaginase* enzyme produced by *E*.*coli*. The bar-chart showed that the intracellular *L-asparaginase* activity was more potential compared to that of extracellular *L-Asparaginase*. Moreover, the *E*.*coli* isolates from cafeteria sewage were more potential *L-asparaginase* producers than those from the River

**Figure 5.**
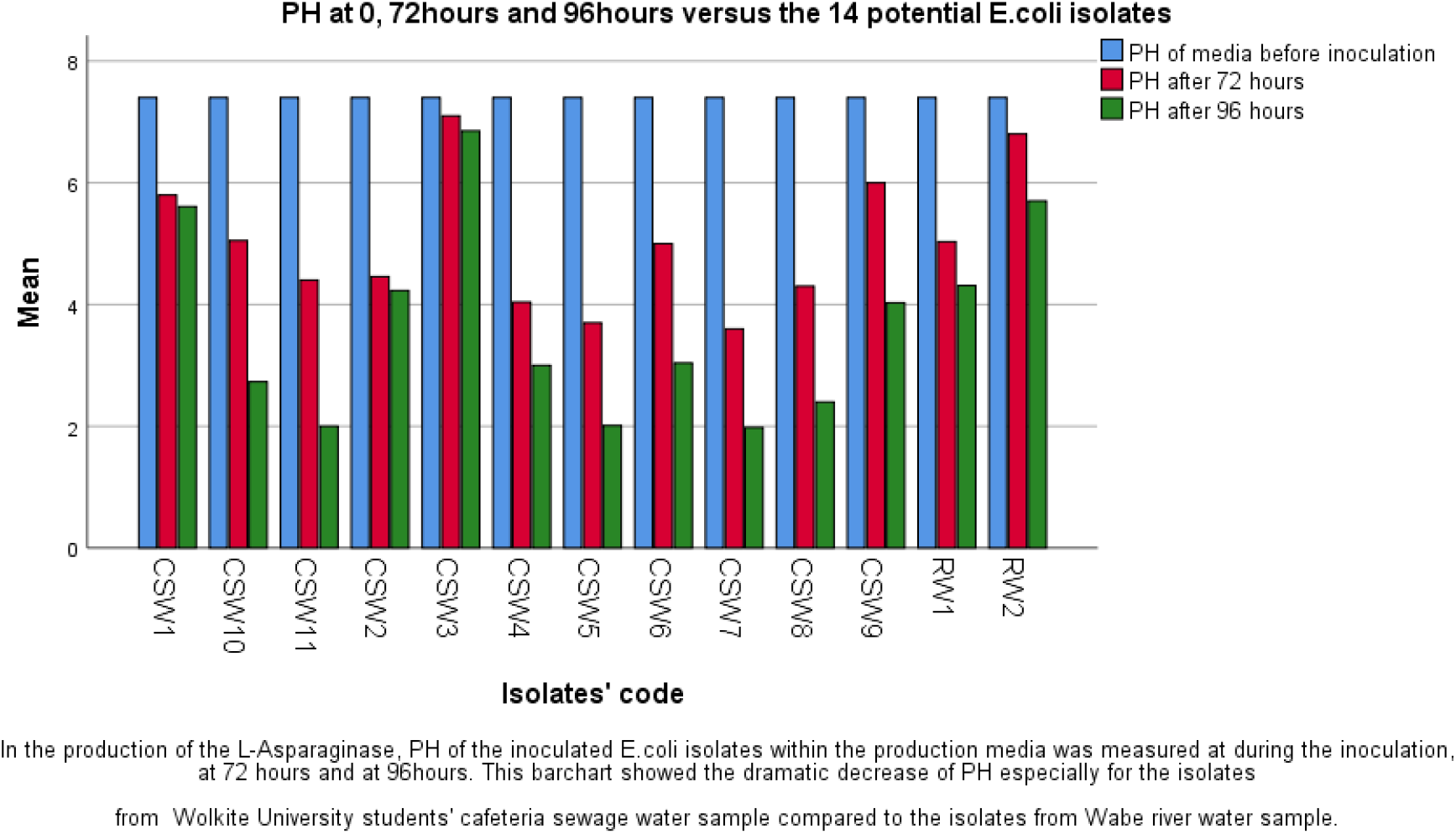
: PH versus the *E*.*coli* isolates: The decreased PH value indicated high production of the *L-asparaginase* enzyme.

### 3.3 Production of L-asparaginase

L-ANase production was carried out by submerged fermentation in 250 mL Erlenmeyer flasks containing 100 mL of sterilized medium. The production media was adjusted at different PH values. A 1ml aliquot of the sample in nutrient broth was inoculated into the medium and incubated in a horizontal shaker at 120 rpm at 37°C for 72 hrs.. A decreased pH indicates that the substrate was digested by the enzyme and separated into acidic monomers, which decreased the PH (Table 1). The bacterial cell mass was separated by centrifugation at 10,000 rpm for 15 min at 4°C. The liquid supernatant was used as an extracellular enzyme. Then, both extracellular enzyme and enzyme within the cell were tested for their activity by measuring the diameter of pinkish red color on the production media. The measure of zone of inhibition for the enzyme within cell was 16.5mm up to 22.25mm while that of extracellular enzyme measured 7.5mm up to 9mm. (Table 1).

**Table 1:** A summary of intracellular and extracellular enzyme zone of inhibition in mm.

| Isolates | Enzyme type | Zone of inhibition in mm | Enzyme type | Zone of inhibition in mm |
| --- | --- | --- | --- | --- |
| CSW1 | Intracellular enzyme | 17 | Extracellular enzyme | 8 |
| CSW2 | Intracellular enzyme | 18.5 | Extracellular enzyme | 7.7 |
| CSW3 | Intracellular enzyme | 16.5 | Extracellular enzyme | 9 |
| CSW4 | Intracellular enzyme | 17.6 | Extracellular enzyme | 8 |
| CSW5 | Intracellular enzyme | 19 | Extracellular enzyme | 8 |
| CSW6 | Intracellular enzyme | 22.25 | Extracellular enzyme | 9 |
| CSW7 | Intracellular enzyme | 22 | Extracellular enzyme | 7.54 |
| CSW8 | Intracellular enzyme | 21 | Extracellular enzyme | 7.5 |
| CSW9 | Intracellular enzyme | 20.5 | Extracellular enzyme | 9 |
| CSW10 | Intracellular enzyme | 21.5 | Extracellular enzyme | 9 |
| CSW11 | Intracellular enzyme | 18 | Extracellular enzyme | 8 |
| RW1 | Intracellular enzyme | 17.7 | Extracellular enzyme | 9 |
| RW2 | Intracellular enzyme | 20 | Extracellular enzyme | 7.6 |
| RW3 | Intracellular enzyme | 22 | Extracellular enzyme | 8.5 |

**Table 2:** Production of *L-Asparaginase*: The PH of the 14 potential *E*.*coli* isolates within the production media was dramatically decreased. The PH measure was conducted at 24 hours time interval.

| Isolates' code | PH of media before inoculation | PH after 72 hrs | PH after 96 hrs |
| --- | --- | --- | --- |
| CSW1 | 7.4 | 5.8 | 5.61 |
| CSW2 | 7.4 | 4.46 | 4.23 |
| CSW3 | 7.4 | 7.1 | 6.85 |
| CSW4 | 7.4 | 4.04 | 3 |
| CSW5 | 7.4 | 3.7 | 2.01 |
| CSW6 | 7.4 | 5 | 3.04 |
| CSW7 | 7.4 | 3.6 | 1.98 |
| CSW8 | 7.4 | 4.3 | 2.4 |
| CSW9 | 7.4 | 6 | 4.03 |
| CSW10 | 7.4 | 5.05 | 2.73 |
| CSW11 | 7.4 | 4.4 | 2 |
| RW1 | 7.4 | 5.03 | 4.31 |
| RW2 | 7.4 | 6.8 | 5.7 |
| RW3 | 7.4 | 6.71 | 5.5 |

### 3.4 PCR confirmation of potential L-Anase producer *E*.*coli* isolates

Ploymerase chain reaction was used to confirm presence of L-Asparaginase protein coding gene named *ansA* gene in the *E*.*coli* isolates (Mohamed *et al*., 2015). Clear, specific bands at the expected size of about 300bp in the sample lanes were observed (Figure 6). The PCR result depicted that the *ansA* gene presence was confirmed in 50% of the isolates.

**Figure 6:**
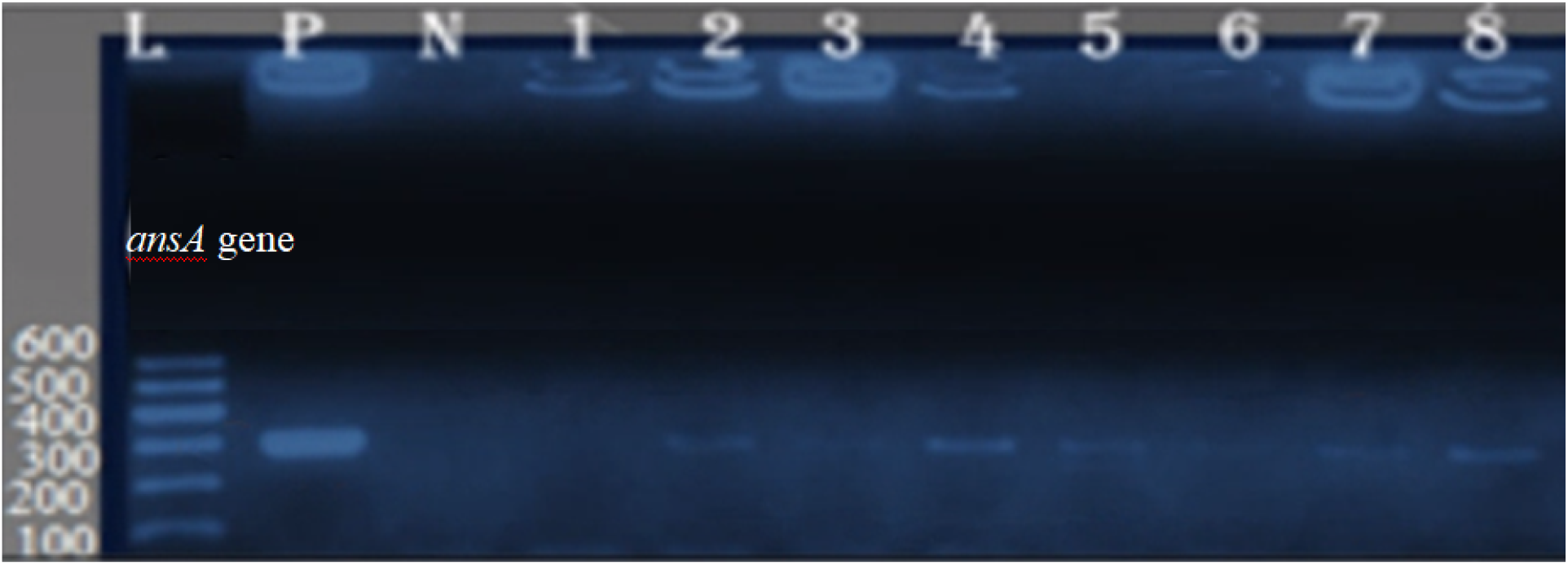
Gel Electrophoresis result of the PCR products

## 4. Discussion

*L-asparaginase* is a promising enzyme with diverse applications, from cancer treatment to food processing. *L-asparaginase* (L-ASNase) is a therapeutic enzyme that has garnered significant attention due to its antineoplastic properties. It catalyzes the hydrolysis of L-asparagine into L-aspartate and ammonia. Its primary applications include treating acute lymphoblastic leukemia (ALL), acute myeloblastic leukemia (AML), and other lymphoid malignancies, often in combination with other drugs. Beyond medicine, L-ASNase is also used in food processing industries to mitigate acrylamide formation (a potential carcinogen) by breaking down L-asparagine. A total of 14 *E*.*coli* isolates were screened for *L-asparaginase* production from the two different sample sites namely Wolkite University students’ cafeteria and Wabe river water sample (Bakeer *et al*., 2022). From the positive isolates those subjected to the screening process, 78.6*%* were from cafeteria sewage water while the rest 21.4% were from Wabe river water sample. Rapid plate assay method was used to test the enzymatic activity by pinkish color development around inoculates on the plate. The pink color depicted the hydrolysis of L-Aspagaine amino acid into aspartic acid and ammonia. Both intracellular and extracellular *L-asparaginase* were checked for their activity on modified M9 media and zone of inhibition was measured for each and every potential isolates (Vimal and Kumar, 2017). The intracellular L-ASNase zone of inhibition ranges from 16.5 to 22.25mm while extracellular L-ASNase measures from 7.5 to 9mm. The result showed that enzyme within the cell showed more inhibition zone than extracellular *L-asparaginase* enzyme. In this process, the *E*.*coli* isolates from cafeteria sewage water still exhibited the largest zone of inhibition which means cafeteria sewage water is the better source of potential L_ASNase producing *E*.*coli* compared to the running river water. A clear, specific band at the expected size in the sample lanes indicates the presence of the ansA gene, suggesting that the isolate may produce *L-asparaginase*. Isolating L-ASNase-producing *E. coli* strains from river ecosystems could provide an alternative and sustainable source. Cafeteria sewage might also harbor L-ASNase-producing bacteria. Cafeteria wastewater contains diverse microbial communities, making it an interesting prospect for screening (Castro *et al*., 2021).

### 5. Conclusion and recommendation

The findings of this study depicted that students’ cafeteria sewage water of Wolkite University is better source for potential *L-asparaginase* enzyme producing *E*.*coli* isolates. Enzymatic activity of the produced *L-asparaginase* was very promising to be used as anticancer drug. Thus, studies investigating the use of these enzymes in search of L-asparaginases, especially those produced through microorganisms, have the capacity to acquire new enzymes with acceptable properties. These discoveries ought to be accompanied through in-depth paintings aiming to increase the process to enable and extend the use of this enzyme, mainly in industrial sectors. Understanding this, molecular biology tools are useful because they indicate the need for further work, including extraordinary method configuration evaluation, in addition to the use of bioreactor options.

## Acknowledgement

We want to extend our immense thank to Wolkite University for allowing us all the laboratory materials needed to conduct the research.

## Source of fund

The research was fully funded by Wolkite University

## Declaration

The authors declared that there is no conflict of interest

## Data Availability Statement

The data that support the findings of this study are available from the corresponding author upon reasonable request.

## Notes

### Competing Interest Statement

The authors have declared no competing interest.

## References

Abdelrazek, N.A. et al. (2020) ‘Production, characterization and bioinformatics analysis of l-asparaginase from a new Stenotrophomonas maltophilia EMCC2297 soil isolate’, AMB Express, 10(1). Available at: 10.1186/s13568-020-01005-7.

Acharya Tankeshwar (2023) ‘EMB Agar Composition, Principle and colony morphology’, microbeOnline [Preprint].

Bakeer, W. et al. (2022) ‘Isolation of asparaginase-producing microorganisms and evaluation of the enzymatic antitumor activity’, Egyptian Pharmaceutical Journal, 21(3), pp. 282–292. Available at: 10.4103/epj.epj_11_22.

Castro, D. et al. (2021) ‘L-asparaginase production review: bioprocess design and biochemical characteristics’, Applied Microbiology and Biotechnology, 105(11), pp. 4515–4534. Available at: 10.1007/s00253-021-11359-y.

Gopireddy, V.R. (2011) ‘Biochemical tests for the identification of bacteria’, International Journal of Pharmacy and Technology, 3(4), pp. 1536–1555.

‘Key facts’ (2018), (February).

Mahajan, R. V. et al. (2013) ‘A rapid, efficient and sensitive plate assay for detection and screening of l-asparaginase-producing microorganisms’, FEMS Microbiology Letters, 341(2), pp. 122–126. Available at: 10.1111/1574-6968.12100.

Mohamed, Z.K. et al. (2015) ‘Cloning and molecular analysis of L-asparaginase II gene (ansB)’, Journal of BioScience & Biotechnology, 4(3), pp. 291–302. Available at: http://search.ebscohost.com/login.aspx?direct=true&db=a9h&AN=116513432&site=ehost-live.

Now, B. (2024) ‘Isolation and identification of Escherichia coli (E. coli)’.

Sharma, S., Verma, R. and Pandey, L.M. (2019) ‘Crude oil degradation and biosurfactant production abilities of isolated Agrobacterium fabrum SLAJ731’, Biocatalysis and Agricultural Biotechnology, 21(June). Available at: 10.1016/j.bcab.2019.101322.

Sudhir, A.P. et al. (2012) ‘Production and amplification of an L-asparaginase gene from actinomycete isolate Streptomyces ABR2’, Annals of Microbiology, 62(4), pp. 1609–1614. Available at: 10.1007/s13213-011-0417-0.

Vaishnavi, L. and Palaniswamy, R. (2023) ‘SOURCES AND APPLICATIONS OF L-ASPARAGINASE: A REVIEW’, 8(12), pp. 396–400.

Vimal, A. and Kumar, A. (2017) ‘Biotechnological production and practical application of L-asparaginase enzyme’, Biotechnology and Genetic Engineering Reviews, 33(1), pp. 40–61. Available at: 10.1080/02648725.2017.1357294.

